# Benchmarking scRNA-seq imputation tools with respect to network inference highlights deficits in performance at high levels of sparsity

**DOI:** 10.1101/2021.04.02.438193

**Authors:** Lisa Maria Steinheuer, Sebastian Canzler, Jörg Hackermüller

## Abstract

Gene correlation network inference from single-cell transcriptomics data potentially allows to gain unprecendented insights into cell type-specific regulatory programs. ScRNA-seq data is severely affected by dropout, which significantly hampers and restrains current downstream analysis. Although newly developed tools are capable to deal with sparse data, no appropriate single-cell network inference workflow has been established. A potential way to end this deadlock is the application of data imputation methods, which already proofed to be useful in specific contexts of single-cell data analysis, e.g., recovering cell clusters. In order to infer cell-type specific networks, two prerequisites must be met: the identification of cluster-specific cell-types and the network inference itself.

Here, we propose a benchmarking framework to investigate both objections. By using suitable reference data with inherent correlation structure, six representative imputation tools and appropriate evaluation measures, we were able to systematically infer the impact of data imputation on network inference. Major network structures were found to be preserved in low dropout data sets. For moderately sparse data sets, DCA was able to recover gene correlation structures, although systematically introducing higher correlation values. No imputation tool was able to recover true signals from high dropout data. However, by using an additional biological data set we could show that cell-cell correlation by means of specific marker gene expression was not compromised through data imputation.

Our analysis showed that network inference is feasible for low and moderately sparse data sets by using the unimputed and DCA-prepared data, respectively. High sparsity data, on the other side, still pose a major problem since current imputation techniques are not able to facilitate network inference. The annotation of cluster-specific cell-types as a prerequisite is not hampered by data imputation but their power to restore the deeply hidden correlation structures is still not sufficient enough.

## 1 Introduction

Inferring gene regulatory networks from transcriptomic data has led to important biological insights for example in recurrence-associated genes of colon cancer [39] or gene architectures associated with Alzheimer’s disease [1].

However, current knowledge is mainly based on information collected from a population of cells which, depending on the experimental setup, might comprise different cell (sub)types, cells resting in different cell cycle stages, or cell subpopulations that gained certain genetic mutations.

With the technological breakthrough of single-cell genomics it is now feasible to extract and analyze the transcriptome of each cell separately, leveraging those obstacles and opening up a novel level of resolution. For example, scRNA-seq helped to identify previously masked cell types [9, 17], to uncover cellular dynamics within a cell population [4], and to unfold developmental trajectories [35, 13]. The inter-cellular heterogeneity in single-cell transcriptomics data furthermore promises to substantially improve our understanding of cellular processes, allowing to uncover gene regulatory networks that are specific to a cell type or cluster of cells and that so far remain buried [31] (see Figure 1).

**Figure 1.**
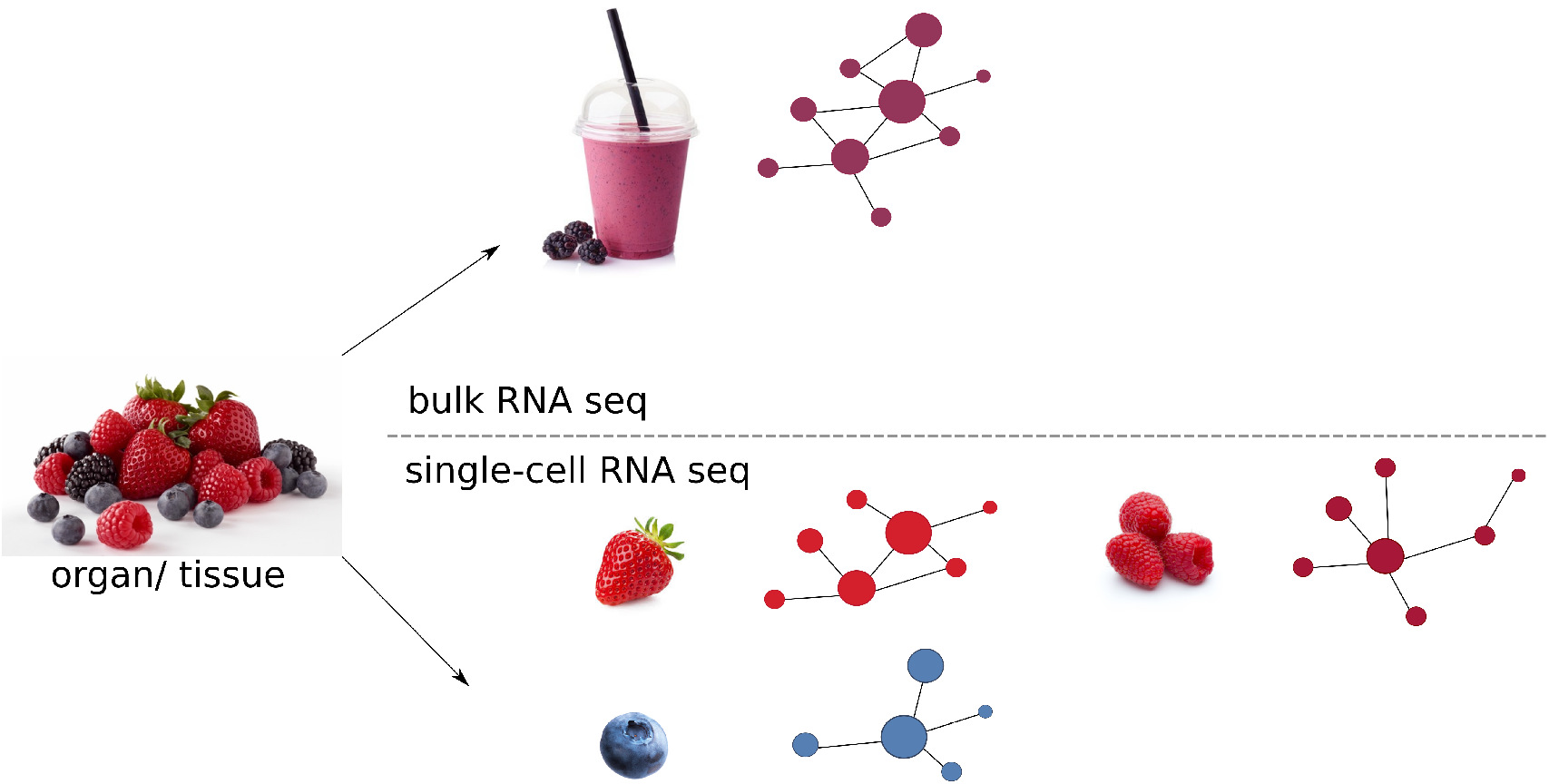
Difference in resolution from bulk and single-cell RNA-seq data on the level of gene networks. Illustrating the input of sequencing experiments as various fruits (left) would result in a homogenized, mixed source material for bulk RNA-seq data, much like a fruit smoothie (top). Inferred networks would therefore reflect an average of all the signals detected in the mixture of, potenially different, cells. Single-cell RNA-seq on the other hand would preserve the information of the individual fruits, hence different cell types and would allow for cell type specific gene correlation networks (bottom). Thus similarities as well as differences in netwoks between cell types could be carved out that would be hidden or covered in the bulk RNA-seq data.

However, to fullfill this promise, two things must be achieved using scRNA-seq data: (*i*) the identification of cell types, and (*ii*) the possibility to infer gene correlation networks.

A plethora of novel analysis tools and comprehensive pipelines has been developed in recent years [34, 5] ^1^ trying to harness the special characteristics of single-cell transcriptomics data. Next to its shear size, the data is extremely sparse and afflicted with two types of zero values [20]. They can either represent a truely absent count or be introduced methodologically where a transcript was expressed but not measured. This second type of zero, referred to as sparsity, can therefore arise from two different scenarios: either systematically, such as mRNA degradation during cell lysis, or by chance where a barely expressed transcript is measured in one cell but not in the other though present in both [18]. The latter phenomenon is also denoted as ‘dropout’. In addition, single-cell data owns higher degrees of variability and technical noise compared to bulk RNA-seq data [8]. Allover, these specific characteristics of scRNA-seq data may interfere with the previously stated aims, particularly regarding network inference from cell-clusters.

While single-cell transcriptomics data sets were restricted to only a few cells, containing far less sparsity, early `WGCNA`-based approaches however allowed new insights into genetic programs [23, 36]. With the emergence of higher throughput methods, these bulk-derived, non-single-cell specific tools showed only mediocre results [8]. Subsequent developments of highly specific single-cell tools, however, were also not able to transform the unique characteristics of this data into the desired outcome [8]. More recent work benchmarked single-cell specific tools using curated reference models [28, 24]. Including 12 algorithms as well as three different sources of reference datasets, `BEELINE`, for example, offers an evaluation framework to facilitate development of novel tools [28]. While testing conditions were improved, network inference remains a challenging problem. Although algorithms such as `SCENIC` or `CellOracle` [2, 16] have been proven to be meaningful, they calculate the network inference on a restricted search space by subsetting the number of investigated genes, and thus still neglect an overall and unbiased picture.

One potential direction to improve the efficacy of single-cell analysis tools is to leverage the data sparsity through imputation approaches intending to ‘interpolate’ the previously missed expression. In the past years, many single-cell specific imputation tools have already been published [15, 11, 27] and their evaluation showed that denoising sparse simulated data can, for example, help to reobtain original cell clusters and time-course patterns [10]. Because of the rapid increase of published imputation methods, several review articles [20, 7] and benchmarking analysis [40, 25] have been published in recent months, also investigating specific fields within the downstream analysis realm, such as differential gene expression [14]. Unsupervised clustering of cells represents another common downstream tool in the analysis of scRNA-seq data which allows for later cell type classification. Past work already proofed that some imputation tools did improve clustering compared to the unimputed situation [14]. Apart from the overall clustering performance, the impact of data imputation on specific cell cluster expression profiles was investigated. Since each cell type owns a set of specific marker genes, data imputation should not render or destroy these expression profiles, as they take a central part in the inference of cell type specific network configurations, but recent research revealed that their reprodicibility was reduced after data imputation [3].

However, the precise influence of data imputation on gene correlation networks remained out of focus. In general, the assessment of cell-type specific network inference based on scRNA-seq data that builds on imputation and evaluates inference and cell type identification cluster-specific expression profiles proofed to be difficult [6]. Since any experimentally derived scRNA-seq dataset is affected by a considerable degree of dropout, a synthetic gold-standard dataset is required as a reference to assess the performance of network inference. To facilitate statistical evaluation, such a dataset should solely contain ‘true zeros’ and fully exclude dropout. However, such a universal, gold-standard reference dataset is so far missing. Those simulated datasets need to fullfill some criteria to be utilized in a benchmarking study. This strongly includes standard research paradigms like reproducibility, scalability, adjustability, and suitability.

Here we provide a comprehensive benchmarking analysis where we elaborate the precise impact of data imputation approaches for a subsequent network inference on single-cell transcriptomics data. In order to infer the influence of imputation methods, we generated a synthetic, non-sparse, ground truth dataset through the downsampling of a bulk RNA-seq dataset, containing solely true zeros and correlation structure, both, between cells and genes, in contrast to other approaches. Based on this dataset, six additional reference datasets were generated with increasing levels of dropout, thus allowing for conclusions on different real-world scenarios. Representative imputation methods have then been applied to all reference datasets and their corresponding network structures were compared to the structure of the gold data.

## 2 Results

### 2.1 Generation of single-cell like reference datasets

In the following paragraphs, we will describe the generation of our reference datasets. In this respect the term gold data refers to the single-cell like data that contains no zeros that are inserted based on sequencing errors (dropout). Alltogether the gold and all six dropout affected datasets are referred to as reference data.

#### Gold data

While scRNA-seq allows for unprecedented biological insights, technical as well as biological noise produce an extremely sparse data matrix hindering current network inference methods [28, 8, 24]. In order to desparsify data, imputation approaches might help to overcome this constraint, thus paving the way for well known network inference tools.

Since real scRNA-seq data always suffer from dropout, we generated a synthetic gold dataset, which only contains true zeros. To ensure reproducibility, scalability, adjustability, and especially suitability of this gold data, an existing mouse hair follicle bulk dataset by Wang *et al.* [33] was used to simulate single-cell data, containing 48,795 genes across 48 conditions (over 20 cell types). For the downsampling procedure described by Peng *et al.* [27], eight hair follicle cell types were chosen randomly such that the synthetic gold dataset owned 5000 genes across 800 cells. After a quality filtering to remove barely expressed genes, being the consequence of the sampling procesure, we finally created a gold dataset with 4960 genes across 800 cells. In total, this dataset contains 15% of true zeros. As a proof-of-concept, we applied `WGCNA` [21] to the gold dataset to ensure that we retained the natural correlation structure of the bulk RNA-seq dataset, see Figure 2. Figure 2A shows the *R*^2^ of a fit to a scale-free topology model, which is a hallmark of many networks, over different soft threshold *β* values. Based on this plot, *β* is chosen to generate a network with approximately scale-free topology. For the gold data a plateau for *β* values larger than eight was reached. Other simulation tools, like `splatter` [38] or `GeneNetWeaver` [30] failed in producing datasets that allowed to reconstruct approximately scale-free networks. This may be due to an insufficient ability of these tools to correspond to real biological processes that has been previously reported [28]. By choosing a *β* of nine for the gene network inference, the gene dendrogram showed a clear hierarchical structure within the genes (Figure 2B). In total, seven different gene modules were detected, whereby the turquoise module was the largest with 2143 genes and the black module was the smallest with only 21 genes.

**Figure 2.**
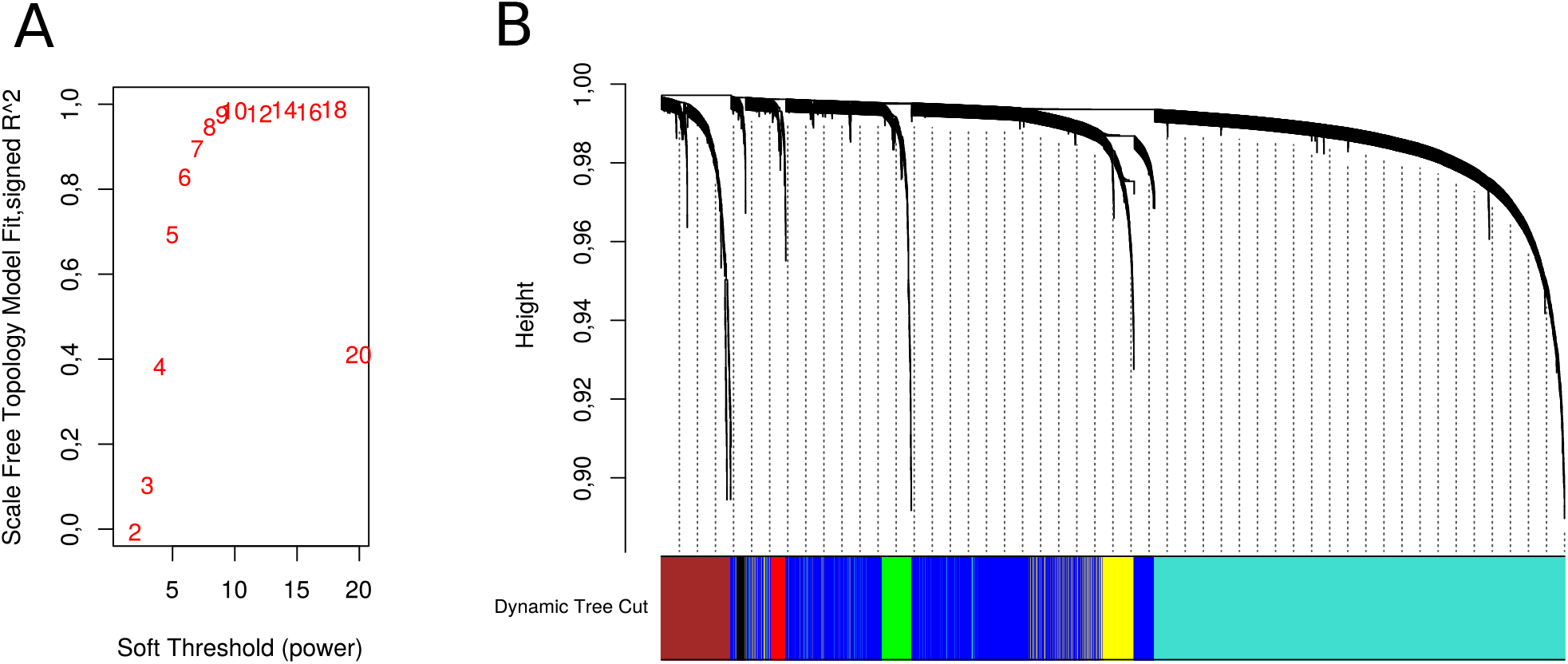
Gold data characteristics. **(A)** Scale-free model fit of gold data using the `WGCNA` package. Development of scale-freeness over twenty *β* values. A plateau of *R*^2^ values was reached for *β* values larger than eight. **(B)** Cluster Dendrogram of gold data using the `WGCNA` package. The gene module assignment is shown in the color bar below. Hierarchy of the genes is indicated on the y-axis. This result is based on a *β* of nine. In total seven gene modules were defined.

#### Reference datasets

Based on the gold dataset, we generated six additional datasets with increasing degrees of dropout, ranging from 40% to 84%. With the 15% true zeros in the gold data, we hence created six datasets containing 55% to 99% percent zeros, which resembles the proportion of zeros of current scRNA-seq methods. The characteristics of these reference data sets are summarized in Figure 3. With an increasing degree of dropout, the distribution of gene expression values indicates the expected shift towards larger proportions of lowly expressed or missing genes (see a clipped version in Figure 3A and the complete graph in Supplemental Figure S4). The gold data shows the smallest fraction of lowly expressed genes, while the 84% dropout dataset shows the opposite behaviour. A real scRNA-seq dataset of human retina organoid cells [19], which was generated via the 10X Genomics procedure was included and plotted for comparison, exhibited a very similar distribution as the 84% dropout dataset (Figure 3A).

**Figure 3.**
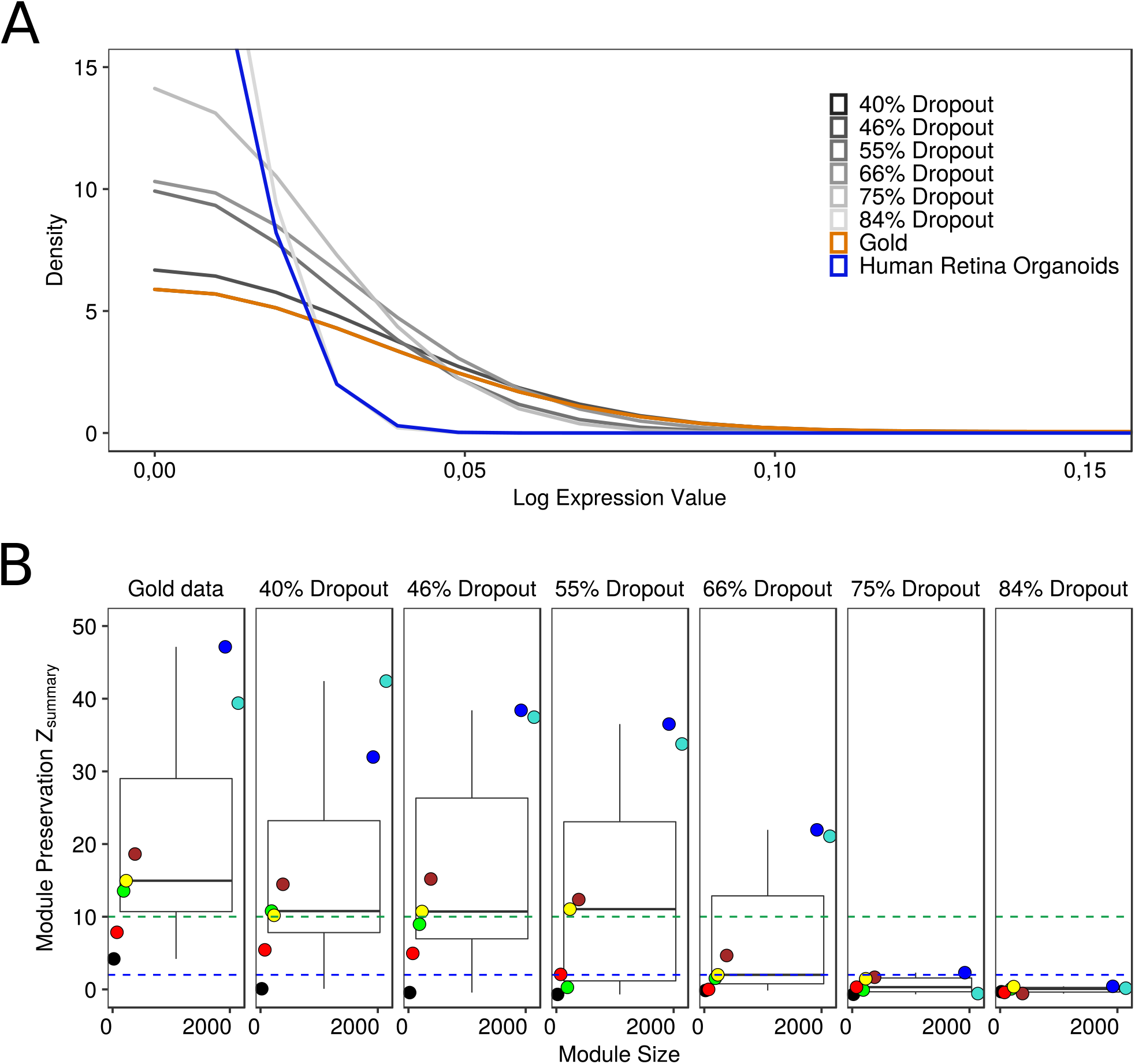
Reference data characteristics. **(A)** Distribution of logged expression values of all reference datasets and the human retina dataset. Density of expression values across eight datasets is shown to contrast impact of different dropout rates. The gold data is plotted in orange, all six dropout reference datasets are shown with a grey gradient and a biological dataset is plotted in blue. Here, only an excerpt between 0.0 and 0.2 is shown. The full plot is provided in Supplemental Figure S4. **(B)** Boxplots showing the behavior of module preservation across different dropout levels. The Z_summary_ measure implemented in `WGCNA` is a composite, permutation-based metric of various network density and connectivity measures. The blue and the green line indicate the threshold towards moderate and strong module preservation, respectively. Coloration of the dots correspond to the individual modules. Dropout refers to the the amount of artificially introduced non-true zeros in each of the reference datasets. An increase in dropout is associated with a decrease in module preservation.

We investigated to what extent hallmarks of network structure can be recovered, or are lost with increasing degrees of dropout. (Figure 3B). For this purpose we utilized the *Z*_summary_ statistic from WGCNA which quantifies module preservation [22] for all reference datasets. This measure indicates how well a group of correlated genes (modules) from a reference dataset is preserved in a test data set. Here, all artificially dropped out datasets were compared to the gold data. Since *Z*_summary_ is dependent on module size, we compared the gold dataset to itself as a reference. As introduced by Langfelder and colleagues, two different thresholds were applied defining the preservation of gene modules [22]. Modules with a *Z*_summary_ value below two are considered to be not preserved at all, whereby values larger than ten indicate a strong preservation. Modules in between both thresholds are considered moderately preserved. Within the gold data, modules showed a median *Z*_summary_ value above the strong preservation threshold as also seen for slightly increased dropout levels (40% – 55%). *Z*_summary_ values close to zero were detected for both high sparsity datasets containing 75% and 84% dropout. Moreover, no single module remained even moderately preserved in the 84% dropout reference dataset. In general, we can state that an increase in dropout correlates with a decrease in module preservation. Nonetheless, correlation network appears fairly robust for low to intermediate levels of dropout. Up to a level of 55 % dropout we observed strong preservation for the majority of modules.

Hence, we were able to generate reference datasets that (1) resemble true biological datasets in their distribution of expression values, (2) retain the natural correlation structure of its original bulk RNA-seq ancestor, and (3) were shown to have a decreasing network preservation with higher levels of dropout.

### 2.2 Impacts of data imputation on network inference

Since dropout seems to compromise network inference we next investigated, whether imputation methods alleviate the situation.

We selected five imputation tools that each represent a certain class of imputation methods. Whereby `DrImpute`[11], `SAVER`[15] and `ENHANCE`[32] are smoothing, model-, or low-rank matrix-based, respectively, `DCA`[10] uses a neural network structure. `scNPF`[37] performs imputation while extracting gene correlation network information from the dataset. All tools are highly used and well-accepted in the single-cell community. A sixth method, `DISC`[12], belongs to the class of neural networks but performs imputation in a semi-supervised manner. For more information, please see the methods section.

#### Module Preservation

To compare the imputation tools on the basis of *Z*_summary_ values, we computed a module-wise *Z*_summary_ log-2 fold change (log2-FC) that reflects the difference in module preservation to the gold dataset before and after imputation, see Figure 4A. A negative fold change therefore represents a module preservation smaller than the gold data and thus a loss of network information.

**Figure 4.**
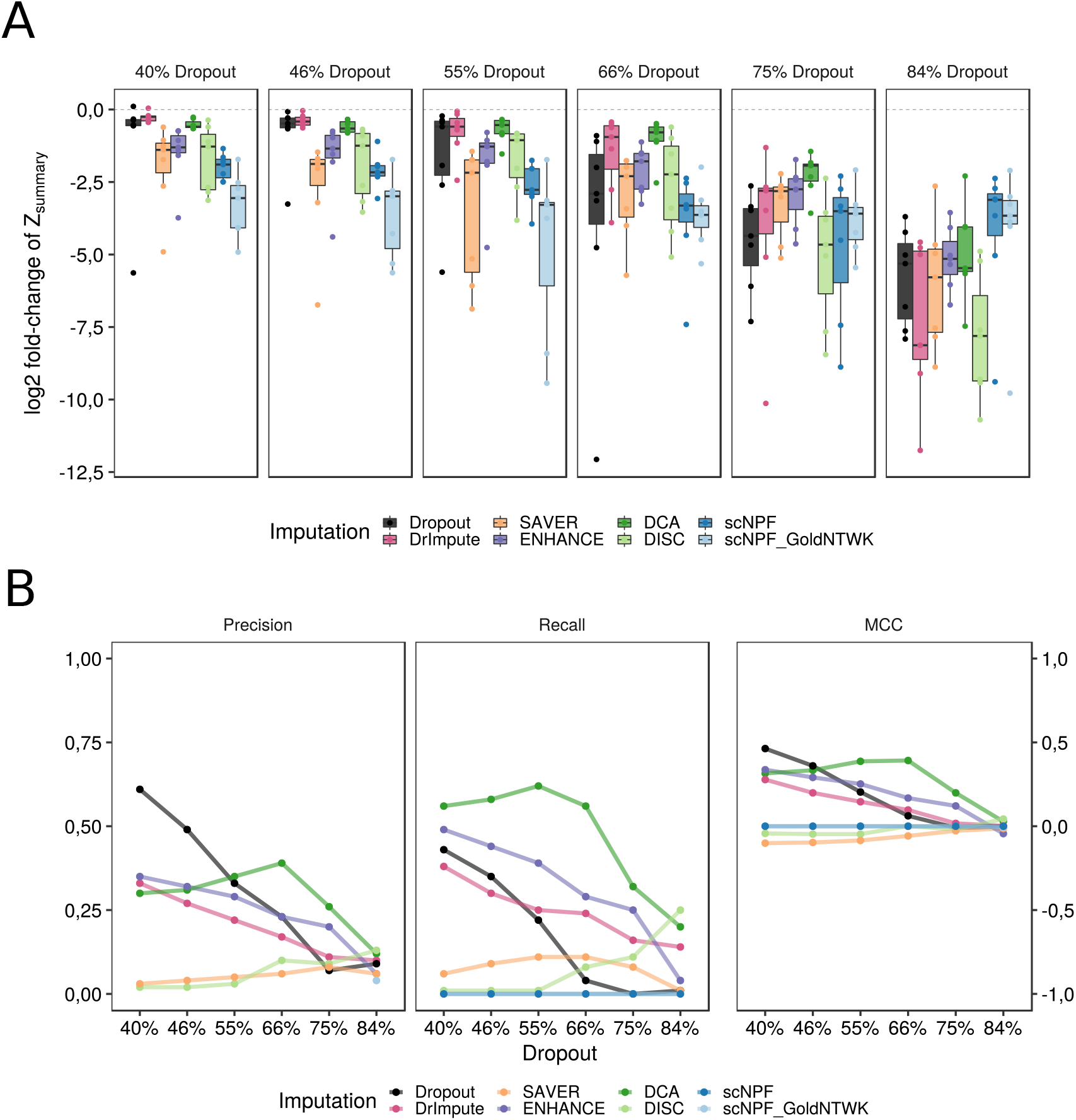
Network measures before and after imputation of reference datasets. **(A)** Boxplot of the *Z_summary_* log2 fold change (log2-FC) of all reference datasets compared to gold data before and after imputation. *Z_summary_* was computed for gold data compared to itself and for any reference data set, with and without imputation, compared to gold data, respectively. Subsequently, a log2 fold change was computed between the gold versus gold and any reference versus gold *Z_summary_* to illustrate how well gene modules from the gold data were preserved in the dropped out and imputed data. The dashed line indicates completely recovered modules. Dots represent values of individual gene modules. The dropout data is depicted in black. Implementation tools that employ related methodological approaches were grouped by similar colors. **(B)** Trends of Precision, Recall, and Matthews correlation coefficient (MCC) over all dropout levels indicating the ability to recover true edges after binarizing the gold data network. Mean and standard deviation (*SD*) of the topological overlap matrix (TOM) per dataset were computed. Edges with a TOM greater 1 * *SD* + *mean* of the gold and all dropped out and imputed datasets were retained. Retained edges of gold data were considered as true edges for computing Precision, Recall, and MCC, respectively.

The unimputed, dropout-affected reference datasets (shown in black) showed lower negative log2-FC values with an increase in dropout, reflecting the loss of module preservation as it was described above. All log2-FC in the 40% dropout dataset were still close to zero, while it decreased drastically in the 84% dropout dataset. If imputation restored the correlation structure, values closer to zero would be obtained. However, we observed a rather diverse picture: For low levels of dropout, where network inference still worked robustly without imputation, only `DrImpute` and `DCA` yielded a log2-FC comparable to the unimputed data. All other tools performed considerably worse. At intermediate levels of dropout (55–66%), where network inference without imputation was increasingly affected but still feasible, `DrImpute` and `DCA` strongly improved log2-FC. `SAVER`, `ENHANCE` and `DISC` resulted in similar log2-FC distributions as the unimputed data, while both variants of `scNPF` even diminished the network inference.

For the high dropout levels, none of the imputation tools enabled network inference with sufficient log2-FC. While `DCA` and `ENHANCE` for 75% dropout and `scNPF` for 84% of dropout performed substantially better than the unimputed data most of the network structure was still lost. An overview of the module preservation values is presented in Supplemental Figure S5.

In summary, from a module preservation perspective, imputation using `DrImpute` and `DCA` facilitated network inference for datasets with low to intermediate levels of dropout, whereas several other tools severely compromised the inference at those dropout levels. For high-level dropout datasets, however, no imputation tool showed convincing and promising results.

#### Edge recovery

By calculating module preservation scores, we employed a similarity measure to infer the impacts of imputation on network inference. By switching the perspective, we additionally wanted to analyze the ability of imputation methods to recover gold data gene interactions.

Therefore, we transformed the network with continuous edge weights to an unweighted network either stating that an edge is present or not. This transformation allowed the computation of measures like precision, recall, and Matthews correlation coefficient (MCC) to quantify the potential to recover true edges. To account for different data distributions and ranges, the binarization threshold was derived individually per dataset. The thresholds were calculated based on the mean and standard deviation of the topological overlap matrix (TOM) with 1 * *SD* + *mean*, meaning that only weights above this threshold were accounted as existing edges (see Figure 4B). Results using 2 * *SD* + *mean* are depicted in Supplemental Figure S2. Initially, we evaluated the precision, which calculates the fraction of true edges over all detected edges, see Figure 4B. The higher the precision, the less false edges were detected. Before imputation, the dropped out reference datasets revealed moderate precision in the 40% and 46% dropout datasets. With increasing dropout, precision dropped towards zero. Imputation of low sparsity datasets (40–55% dropout) did not improve the precision. Whereby some tools such as `DCA`, `DrImpute`, and `ENHANCE` reached precision values half as high as the dropped out data, `SAVER`, and `DISC` revealed values close to zero. We could not calculate precision values for both `scNPF` approaches. Based on a high mean and a high standard deviation in the TOM values, we did not retain any edge. As soon as the dropout level exceeded 55 %, `DCA` retrieved a higher precision than the unimputed data with a peak of performance at the 66% dropout dataset. For both high dropout datasets, all approaches showed low precision values. These results suggest, that independent of the method used, imputation methods applied to high dropout datasets tend to inflict the correlation network with large amounts of false edges.

The recall measures the fraction of true edges in the gold network that were actually classified as edges in the dropout or imputed datasets. Similiar to the precision, the dropout datasets revealed a stepwise decrease in recall with increasing dropout level. Overall, this trend also persisted after data imputation. However, `DCA` showed increasing recall rates until a dropout level of 55% is reached. `DCA`, `ENHANCE`, and `DrImpute` consistently outperformed the unimputed datasets. `DISC` showed a steady increase in recall starting from dropout levels of 55%, resulting in the highest recall value at 84%. Both `scNPF` resulted in recall rates of zero - based on the fact that no edge was detected in those datasets.

Similar to scalefree biological networks with only a few highly connectes edges, the datast was highly unbalanced towards true negativ edges. The MCC represents a more robust evaluation of the classification for such inbalanced datasets since it makes use of all entries in the confusion matrix. Perfectly predicted values reflect an MCC of one, whereby a random assignment would result in values close to zero. Predictions opposite to the true values have negative values ranging up to minus one. Here, we observed trends similar to precision. While the dropout datasets exhibited the highest MCC values at 40 and 46% dropout, `DCA` and later `ENHANCE` outperformed the unimputed data exceeding 46% dropout. For the highest dropout dataset, neither the sparse nor any imputation tool reached good MCC values. Both `scNPF` approaches and `SAVER` consistently showed values close to or below zero.

Gaining a broader picture of the effect of imputation prior to network inference, the edge recovery analysis revealed that good recall values in data imputation are mainly caused by inflating the data with more correlation, i.e. artificial edges, compared to the Gold data as highlighted by the moderate precision values (Supplementary Figure S6). However, as already indicated by the module preservation analysis, overall good edge recovery values were obtained for the unimputed data with low dropout levels. `DCA` revealed a moderate performance on datasets affected with intermediate levels of dropout.

### 2.3 Cell cluster annotation in human retina organoid data

Apart from gene network inference, we also investigated cell correlation before and after imputation, which are essential to annoate cell types prior to identifying cell-type-specific network properties. Therefore we used a human retina organoid dataset that was published by Kim *et. al* [19]. This dataset is afflicted by dropout (total percentage of zeros: 85%) and will be denoted hereafter as sparse dataset. In their study, Kim *et. al* clustered the cells and annotated them as either rods, cones, or Müller Glia (MG) cells based on (well-established) marker genes extracted from their low-dimensional t- distributed stochastic neighbor embedding (t-SNE). Starting from this sparse dataset we investigated the impact of the previously introduced imputation methods with respect to cell clustering and a subsequent marker gene-based annotation of cell types.

The Louvain cluster detection within the t-SNE of the original sparse dataset revealed eight cell clusters, see Figure 5(a). Most imputation tools produced comparable quantities of eight to eleven cluster, Figure 5(b-f). Solely the imputation by `ENHANCE` resulted in a total of twenty-one clusters being detected (Figure 5(g)). Cluster sizes were comparable between the sparse data and most imputation methods. The largest Louvain cluster contained around 300 cells in the original organoid data, as well as for both `scNPF` variants, `DCA`, and `SAVER`. Markably smaller sizes were obtained after `ENHANCE` and `DrImpute` imputation with 128 and 195, respectively. The smallest Louvain cluster counted between 39 to 46 cells, which was found in the sparse dataset and after both `scNPF`, `SAVER` and `DrImpute`. Deviations were detected after `ENHANCE` and `DCA` imputation with 19 and 62, respectively.

**Figure 5.**
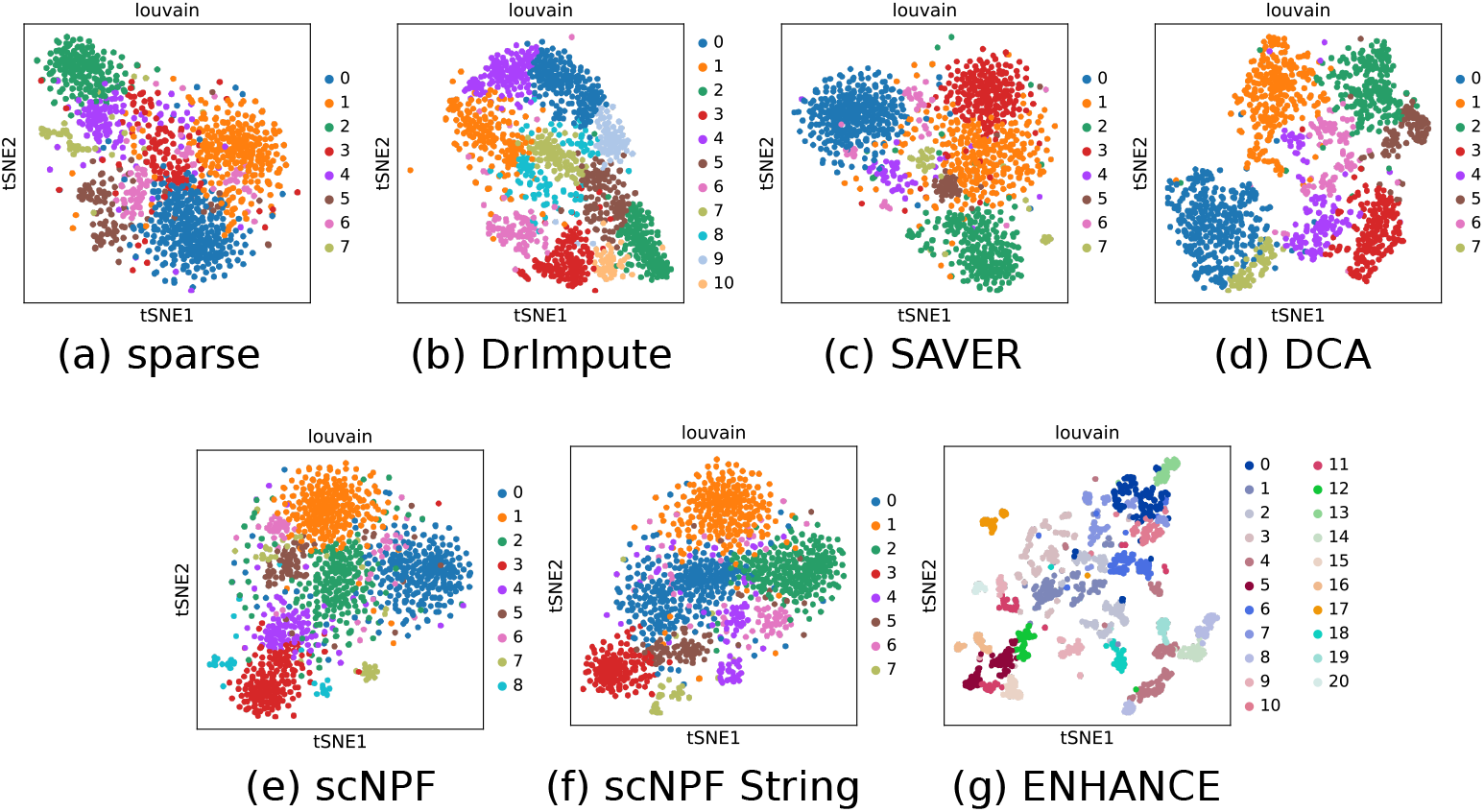
Louvain clusters detected in human retina organoid data before and after imputation. The sparse and imputed datasets were subjected to a standard preprocessing procedure in `scanpy` and projected into a low dimensional t-SNE. The embeddings were colored according to the Louvain cluster detection. Colors are specific for the individual plot and do not correspond between data sets. Eight to eleven Louvain clusters were generally detected. Solely `ENHANCE` imputation resulted in twenty-one clusters. In most cases, cell clusters were found to be overlapping on the cluster boundaries. Visually more separated clusters were obtained after `DCA` and `DrImpute`.

By mere visual inspection of the cluster density for the sparse data, both `scNPF` alternatives, and `SAVER` showed clusters with interwining boundaries. Imputation by means of `DrImpute` and `DCA` resulted in slightly more dense clusters while `ENHANCE` led to a completely different cluster layout with much more fine grained clusters. To back these findings up with a statistical measure, we calculated the mean silhuette coeffiecient (MSC) that allows to quantify the density of clusters. Clusters revealing an MSC of close to one reflect well separated, dense cell clusters, whereas values close to zero hint towards overlapping clusters. Negative values (lowest:-1) indicate that a cell was most likely missclassified. Before and after imputation overall MSC values were found to be close to zero. Slightly positive values were obtained by `DCA` and `ENHANCE`. The weakest performance was found for `DrImpute`. These results support the visual observation of overlapping and not well separated clusters. Thus, data imputation did not improve the cluster density substantially compared to the sparse data.

Based on their original clustering, Kim *et al.*, derived unique marker genes for the retina-specific cell types rods, cones, and MG cells. In the following, the reproducibity and consistency of these marker genes before and after imputation was investigated by comparing quantities of annotated cell types. We set up an automated cluster annotation pipeline based on the expression pattern of these marker gene lists. All dotplots, showing the expression pattern of these marker genes before and after imputation are provided in Supplemental Figure S1. Briefly, a cluster was annotated as rods, cones, or MG cells when more than 75% of the cells in this cluster expressed the respective marker genes. A summary of this analysis is depicted in Figure 6 (bottom). Comparing the amounts of annotated retina-specific cell types before and after imputation, overall similar quantities of rods were detected. More diverging results were found for MG cells. The biggest difference was observed for cones after `ENHANCE` imputation.

**Figure 6.**
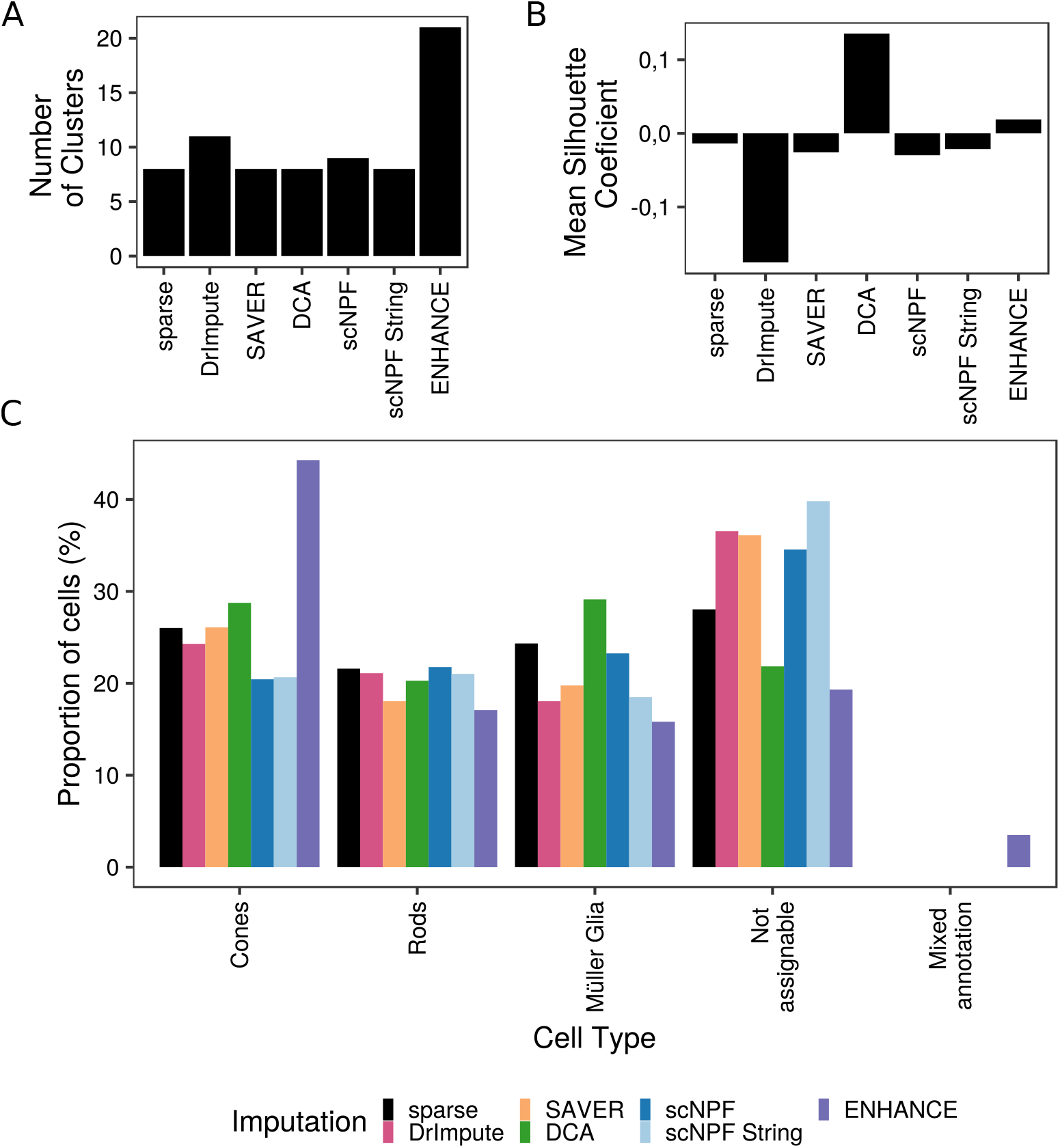
Effect of imputation on cell cluster annotation in human retina organoid data. (top left) The barplot shows the number of Louvain clusters for sparse and imputed data, obtained as described for Figure 5. (top right) The barplot shows the mean silhouette coefficient indicating the overall cluster density. (bottom) Result of the automated cell cluster annotation procedure based on the expression of known marker genes. Expression values per gene and cluster were scaled between zero and one, and binarized (threshold: 0.5). When 75% of the marker genes representing a certain cell type were expressed in a cluster, this cluster was annotated to the respective cell type. The barplot visualizes the percentage of cells annotated as rods, cones, or Müller Glia (MG) cells as well as the percentage of clusters with a mixed, or without a cell type-specific annotation.

To compare the results on a cluster level we defined three additional classes: (a) *pure clusters* represent the percentage of clusters where all cells have the exact same annotation, (b) *mixed annotation* summarize clusters with at least two different annotations, while (c) *not assignable* contain cluster where no unique annotation was possible. Whereby the amount of class (c) was quite different between all imputed and the unimputed data, `ENHANCE` resulted in the lowest and `scNPF String` in the highest pecentages. `ENHANCE` was the only tool that led to cells with mixed annotations (3.5% of cells) (Figure 6C, Supplemental Table S1).

Summarizing the above described results, cluster quantities, clustering performance as well as cluster annotatability behaved comparably before and after imputation. Again, `DCA` improved the results marginally regarding cluster density and the amount of annotated retina cell types. `ENHANCE`, on the other hand, produced a slightly different ratio in cones towards rods and MG cells.

## 3 Discussion

Single-cell omics approaches may provide a unique opportunity to gain unprecendented insights into cell type-specific regulatory programs via the inference of cell-type specific networks and the comparative analysis of these networks between cell types or states. This manuscript investigates to what extent the sparsity or drop-out observed in scRNA-seq data interferes with correlation network inference and the identifcation of cell types using marker genes and whether imputation approaches improve these tasks.

### 3.1 Correlation structure from bulk sequencing data can be used to study the effect of dropout on gene networks in single-cell transcriptomics data

Inferring the effect of dropout on network inference in single cell data requires reference datasets with defined levels of dropout. Since all experimentally generated datasets are afflicted with certain degrees of dropout, this goal can only be achieved with a synthetic dataset. Though several single-cell data simulation tools were available at the beginning of the study, none of those were specifically designed to deliver data with a proper gene correlation structure, which is a trivial requirement of correlation network inference. We therefore applied a downsampling approach to a bulk RNA-seq data set [27] to generate a non-sparse single-cell like gold dataset. We were able to show that this synthetic data has suitable gene-gene correlation properties that enabled the inference of networks of approximately scale-free topology. Since this dataset contains per definition only true zeros we were able to subsequently produce datasets of defined degree of dropout. These reference data sets enabled a thorough investigation of the impact of data dropout on uncovering network information measured as the module preservation compared to the gold network. Although this *Z*_summary_ statistic is dependent on module size, i.e., that smaller modules will decrease faster, we still observed a negative correlation between dropout and module preservation. A similar trend was stated by Zhang *et. al*, where the sum of squared errors increased while Pearson’s correlation coefficient decreases with higher ratios of dropout [40].

We could, however, demonstrate that retrieving reliable and meaningful biological gene networks from low dropout scRNA-seq data is still maintainable (up to 55% dropout). Datasets with dropout levels beyond 75% or even 84%, which resemble, e.g., typical up-to-date 10X Genomics data output, are fairly inappropriate for network inference analysis. An option to overcome this situation could be to manipulate the data prior to the network inference, for example through data imputation. This might help to potentially lift the aforementioned restrictions to finally take advantage of the higher resolution of the source data.

### 3.2 Current imputation methods do not improve network inference from high dropout datasets

Diving deeper into the potentials of imputation approaches, we utilized seven different methods to preprocess our six dropout datasets. By calculating the log2-FC of the *Z_summary_* values compared to the gold data, we were able to investigate the question if those methods allowed to regain at least parts of the burried gene correlation structure, and hence enabled the usage of scRNA-seq data for (sub)network inference.

In general, we observed that the impact of imputation was highly dependent on the applied method and even more so on the dropout level of the source datasets. In addition, our results suggets that most algorithms alter the complete gene correlation structure instead of restoring previously hidden information (see Supplementary Figure S6).

Most imputation tools did not improve module preservation especially for low dropout levels (up to 46%) with the exceptions `DCA` and `DrImpute` which achieved comparable results. However, for such low dropout levels even `DCA` and `DrImpute` fell short of unimputed data in terms of precision of edge recovery suggesting that low dropout datasets barely benefit from imputation. Platforms such as Smart-seq2 are only moderately afflicted by dropout [41]. Hence we would suggest to stick to the original data instead of applying data imputation for such low-dropout data. For intermediate levels of dropout (up to 66%), imputed datasets revealed better module preservation compared to the sparse data. While still `DCA` and `DrImpute` appear to preserve modules best, the highest precision and recall rates of `DCA` compared to all competitors highlighted that `DCA` might be the most suitable option to recover hidden gene correlations in moderately sparse datasets. Beyond 75% of dropout none of the imputation tools was able to approximately restore the true gene correlation structure.

Taking into consideration the results of module preservation and edge recovery, we thus propose to infer networks directly from low dropout single-cell transcriptomics data and use `DCA`-imputed data for intermediate levels of dropout. Although Andrews and Hemberg declare `SAVER` as the ‘safest’ option for data imputation [3], we cannot support this statement based on our findings. In case of high dropout levels, the value and benefit of network inference should be scrutinized in the current situation.

### 3.3 Marker gene expression is not affected by imputation

Next to reliable network inference, identification of specific cellular populations via cell clustering and annotation of clusters is the other important requirement for studying cell type specific networks. One frequently employed approach for cluster annotation is relying on unique expression patterns of known marker genes. After finding that imputation methods tend to induce false positive signals into (high dropout) gene-gene correlation data, we additionally investigated the effect on correlation between cells. Intrinsically, the unique expression profile should prevent marker genes from being too prone to dropout, which makes an imputation not necessarily required prior to cell type annotation. However, in case imputation tools have been applied, it is of major importance that the imputation methods have no negative effect on the overall expression profiles of marker genes. Andrews and Hemberg already pointed out that the reproducibility of marker genes tends to be reduced upon data imputation [3] when extracting and comparing marker genes before and after imputation. Here, we aimed at investigating the impact of imputation on known marker genes more closely employing an experimentally generated dataset.

We obtained clusters without clear distinctions between one another both for sparse data as well as most imputation tools, with the exception of minor improvements for `DCA`. These findings suggest that imputation did not strongly improve cell type separation in low dimensional space. This is in contrast to previous findings by Eraslan `et al` [10], who, however, used a synthetic dataset with lower levels of dropout, which may explain the discrepancy.

Based on the tSNE and Louvain cluster detection, we created an automated annotation pipeline to compare quantities of annotated retina-specific cell types before and after imputation. Allover, annotation of pure retina clusters was only marginally improved using `DCA` or `ENHANCE`. All remaining tools resulted in less retina specific cell types compared to the unimputed data, raising concerns about their applicability and usefulness in this respect. Looking at this analysis part alone, the results suggest that `DCA` as well as `ENHANCE` allow to annotate marginally more retina cells and can thus improve, for example, cell type quantification.

In general, we could show that the applied imputation tools maintain usability of marker genes leading to comparable quantities of annotated cell clusters. Based on this analysis, again `DCA` was able to slightly optimize both, cell clustering and annotatability compared to the sparse situation. Here, no tool led to an eradication of the marker gene profiles, but they have not been found to add certain definiteness to the problem.

## 4 Conclusion

In this manuscript we investigated the influence of scRNA-seq data imputation on gene correlation network inference and annotation of cell clusters. Based on our findings on the preservation of network features in synthetic datasets we recommend to use unimputed data for low levels of dropout and `DCA`-imputed data for datasets with intermediate levels of dropout. None of the investigated imputation approaches allowed to reliably reconstruct correlation networks for high levels of dropout. Cell cluster annotation was neither pronouncedly hampered nor facilitated by imputation. Solely `DCA` slighty improved the clustering of cells. We thus showed that the identification of cell-type specific network properties from scRNA-seq data seems technically feasibly, if dropout is sufficiently well controlled. It remains to be shown that this enables the generation of novel biological insights.

## 5 Methods

In this paper, we investigated the influence of imputation methods with respect to network inference on single-cell RNA-seq data. A visual summary of our analysis workflow is depicted in Figure 7.

**Figure 7.**
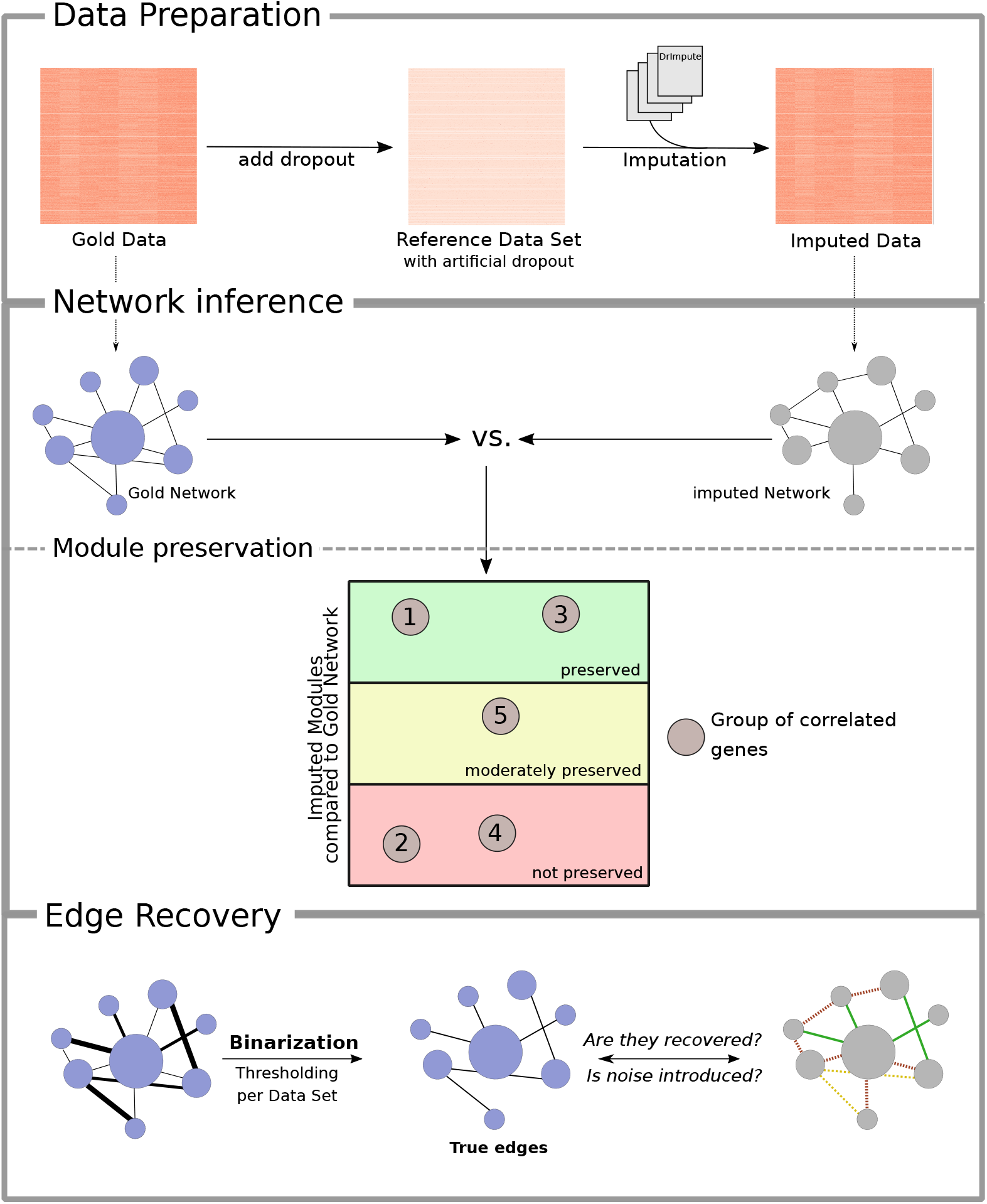
Workflow to investigate the impact of imputation of GRN in single-cell transcriptomic data. By downsampling a bulk RNA seq dataset, a dropout-free, single-cell like, gold dataset was generated. Introducing increasing level of dropout, six artifically sparse datasets were generated which severed as the input for six imputation tools. The ability to preserve correlated gene groups from the gold data was inferred using the quantitativ measure of module preservation from the `WGCNA` package which relies on various network density and connectivity metrics. Apart from the obtained scores, gene modules can be classified as beeing strongly, moderately or not preserved compared to the gold data. In a second step, the ability to recover true edges was evaluated by binarizing the weighted networks. The applied tresholds were determined statistically per dataset individually. Edge recovery was quantified via precision, recall and Matthews correlation coefficient.

The code for data generation, data analysis, and visualization, is publicly accessible on our Github page^2^.

### 5.1 Generation of synthetic reference datasets

To evaluate the ability of reconstructing gene regulatory networks from single-cell RNA-seq data we firstly needed to generate a dropout-free, gold dataset that had to contain an appropriate correlation structure. As shown by Peng *et al.* [27], the existing correlation structure from a bulk RNA-seq datasets can be used to create single-cell-like data. The bulk dataset of mice hair follicles was taken from Wang *et al.* [33]^3^ and contained 48795 genes across 48 conditions. Single-cell data generation was done in accordance to the workflow by Peng *et al.* [27] and is available on their Github page. Briefly, eight conditions and 5000 genes were randomly selected during the downsampling procedure. Each of these conditions was used to simulate a certain cell type. After replicating each cell type 100 times, we generated a 800 cell data set. To resemble the single-cell-like gold data, we used (five times) the standard deviation of each gene in each condition to introduce noise via a random normal distribution with mean zero and one hundred observations. A dropout rate was modelled via the *λ*parameter (range from 0 to 1). The higher the *λ*, the smaller the resulting sparsity. This dropout rate followed an exponential function *e*^−*λ*\*mean expression^2^^. Zero values representing the explicitly dropped out gene expressions, were in the end introduced via a Bernoulli distribution defined by the dropout rate. In total, six different *λ* values were used with 0.01, 0.09, 0.21, 0.42, 0.7, and 0.99 to generate six different sparse datasets ranging from 84 % to 40% dropout, respectively.

All data sets were filtered for genes and samples with missing values using the `WGCNA GoodSamplesGenes` function and transformed to count values.

### 5.2 Imputation methods

This paper aims to investigate the influence of imputation with respect to network inference. To perform a systematic evaluation, we utilized well-established and highly representative tools in accordance to the four classes defined by Lähnemann *et al.* [20]: (1) deep-learning based - `DCA`(`scanpy` version 1.3.1), (2) smoothing-based - `DrImpute`(version 1.0), (3) model-based - `SAVER`(version 1.1.2), and (4) low-rank matrix based - `ENHANCE`^4^. We furthermore created an additonal class (5) of tools which utilize gene networks - `scNPF`(version 0.1.0). In addition to those tools, we also included `DISC`(version 1.1.2), which was very recently published. It also implements a deep-learning approach but performs imputation in a semi-supervised manner. An overview of these tools is given in Table 1.

**Table 1.** Overview of published imputation approaches use.

| Imputation | Class | Data input | Output | Code |
| --- | --- | --- | --- | --- |
| **DrImpute**[11] | (2) | Log10(X+1) data | Logged data | R |
| **SAVER**[15] | (3) | Count data | Count data | R |
| **DCA** [10] | (1) | Count data | Count data | python |
| **ENHANCE**[32] | (4) | Count data | Both possible (Logged used) | R |
| **scNPF**[37] | (5) | Count data + TOM data | Count data | R |
| **DISC**[12] | (1) | Count data | Count data | python |

All imputation tools were run according to their recommended settings which are described in R-markdown files. Solely `ENHANCE` was applied with a fixed knn-parameter of eight for the synthetic dataset. Our reference datasets were provided to these approaches as described in their respective publications. The `scNPF` tool made use of a gene correlation network to guide the imputation process. In its implementation, two different types of networks can be imported which are either derived from the sparse data or extracted from a reference database. Here, we used `WGCNA`(version 1.69) for both scenarios: a network based on the sparse datasets to reflect the first case and a network of the gold data to resemble the latter case. In either way, topological overlap measure (TOM) matrices were used as `scNPF` input. `DISC` required the input data in a `LoomPy`^5^ format which was accomplished in accordance to the tutorial^6^

### 5.3 Network Analysis using WGNCA

After defining the reference datasets as well as the included imputation approaches, the effect on the gene networks must be determined. Thus, gene correlation networks were detected using the Weighted Gene Network Correlation Analysis (`WGCNA`) R-package [21] (version 1.69). The general steps were performed according to the tutorial stated on the website ^7^.

#### Detection of gene clusters

Gene module detection was only performed on the gold dataset which does not contain artificial dropout. After removing genes and cells containing zero values by means of the `GoodSamplesGenes` function, the scale-free model fit was calculated. Here, we achieved a scale-freeness by using a *β* of nine. For subsequent module detection a minimal cluster size of twenty was used while the remaining parameters were left with their default values. The vector including the gene module assignment together with the expression data was used for the calculation of module preservation.

#### Investigate Module preservation statistics

The module preservation was calculated using the provided `modulePreservation` function. Here, a pairwise comparison in module preservation between the gold data and the test datasets (either dropout or imputed data) was calculated. To reduce random reduction of modules, the maximal module size was set to the number of genes included in the dataset. The number of permutations was set to 100, the `dataIsExpr` parameter was left on default, and all other arguments were chosen according to the tutorial. In order to make the results more tangible, log2 fold changes (log2-FC) of the module-specific *Z*_summary_ scores were calculated between the gold and the test datasets. Gold data *Z*_summary_ values were obtained by calculating the module preservation against itself. A negative value indicates a lower *Z*_summary_ value in the test than in the gold dataset. A value of zero corresponds to equal values.

### 5.4 Evaluation of edge detection

Next to module preservation other measures were included to assess whether true gene correlation can be inferred after imputation. We therefore needed to transform the weighted `WGCNA` network to a unweighted, binarized network.

#### Binarization of weighted gene networks

Gene modules were detected in `WGCNA` using the topological overlap matrix (TOM) measure, which does not only account for co-expression similarity, but also for network topology information [21]. By setting certain thresholds to the TOM values, we could binarize the presence of an edge. Since the distribution of TOM values were different before and after imputation, we decided to use a distribution-depended threshold by using the following formular: *e * SD* + *mean*; *e* = {1, 2}. To avoid bias in the model evaluation measures, the self-correlated gene diagonal was removed from the subsequent analysis.

#### Measuring edge recovery

We used the gold network as *true* reference and the imputed networks as *predictions*. Three different measures were applied to analyse the problem from different perspectives. On the one hand, precision and recall were used to approximate the recovery of true gene-correlations. On the other hand, *Matthew’s correlation coefficient* (MCC) was applied to infer the performance of the whole edge classification task. The MCC is particularly useful since these datasets are heavily unbalanced towards true negative entries (non-correlated genes). The measures are defined as follows:

Precision:

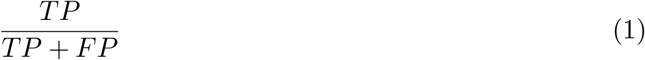

Recall:

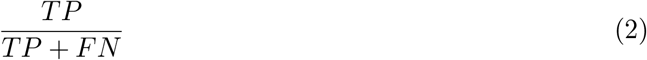

Matthews Correlation Coefficient:

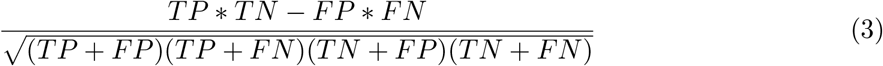

### 5.5 Retina Organoid Dataset

A biological dataset was utilized to evaluate the impact of imputation approaches on the expression profiles of marker genes.

#### Data preparation

We used the human retina organoid dataset from Kim *et. al* (GEO:GSE119343) [19]. In their original study, retina organoids were clustered and annotated according to the major cell types in the retina: rods, cones, and Müller Glia cells. Specific marker genes were extracted to describe each retinal cell type. This dataset contained 19426 genes across 1346 cells and included 85% zeros. Prior to the imputation, genes which were not expressed in at least two cells were removed using the `preprocessSC` function from the `DrImpute` package [11]. Data input was performed as depicted in Table 1

#### Data imputation

`DrImpute, SAVER, DCA, scNPF, DISC`, and `ENHANCE` were applied on the retina dataset. However, `DISC` failed to operate on this large dataset and was therefore not included in this analysis part. All imputation tools were applied using their default parameters as indicated in the corresponding script. Again, `scNPF` was applied using both settings, (1) by extracting a network from the sparse data using `WGCNA` and (2) by incorporating a reference network from the STRING database which was implemented in the tool.

#### Data processing and visualization

In the original publication, the single-cell data was processed with the `Seurat`(version 2.3.4) workflow. In this publication we utilized the `scanpy` pipeline (version 1.6.0)[34] since it proved to be more robust, and adjustable for the needs of this analysis than `Seurat`. Both workflows revealed comparable results, which can be seen in Supplementary Figure S3.

Briefly, sparse and imputated data were filtered for low quality cells and genes upon the initial data import into `scanpy`. As described in the original publication, cells with less than 600 genes and genes which do not occur in at least 5% cells were discarded. Here, 5% account for 67 cells. This parameter was changed for `ENHANCE` imputed data, where genes without any count were removed. Otherwise the subsequent downstream tasks would not have been conductable after imputation.

In a following step, cells expressing an aberrantly high number of genes will be detected and removed. Since some imputation tools introduced a uniform number of expressed genes acrosss cells and to ensure comparability across imputation approaches, this threshold was set individually per dataset to avoid excessive filtering. The individual thresholds are given in Table 2.

**Table 2.** Overview of filtering thresholds used for data preprocessing. The table shows the individual gene number upon which cells were discarded in case they exceed this cutoff of expressed genes.

| **Imputation** | **Number of genes** |
| --- | --- |
| Sparse | 4000 |
| DrImpute | 7900 |
| SAVER | 13369 |
| DCA | 14845 |
| ENHANCE | 14000 |
| scNPF | 13369 |
| scNPF String-reference | 13369 |

Using the data normalization procedure described in the tutorial, followed by a log10(x+1) transformation, highly variable genes were detected and kept using default parameters. Subsequently, unwanted variation from the total gene counts was regressed out and data was scaled according to parameters given in the paper.

We then applied a dimensionally reduction using principal component analysis (PCA) prior to the t-SNE. The neighbor graph was calculated using ten neighbors and 40 principal components. Cluster detection was done via the Louvain algorithm using standard parameters.

#### Clustering performance evaluation

To infer performance of the Louvain clustering before and after the imputation, the mean silhouette coefficient from the `sk.learn` package (version 0.23.2) was used [26]. This score helps to infer clustering performance by investigating the tightness and separation of each cluster [29]. Therein, we utilized the expression data and the Louvain cluster assignment as input. As a metric we used the euclidean distance measure.

#### Automatic cell cluster Annotation

Based on marker genes expression, cell clusters could be annotated to their corresponding cell type. To infer whether this was still possible after the imputation, an automatated cluster annotation pipeline was implemented.

Using the originally published marker genes for cones, rods, and Müller Glia cells [19], the mean expression per gene and Louvain cluster was calculated. To allow for a fair comparison, the data was scaled per Louvain cluster with a min-max-normalization. In a next step, the scaled data was binarized using a threshold of 0.5. A Louvain cluster was then annotated if more than 75% of the marker genes were expressed above this threshold. Three classes of annotation results were inferred such as pure, mixed and unassigned clusters. Pure clusters showed an assignment for exactly one cell type and mixed clusters exhibited at least a double assignment. The third class included Louvain clusters with no successful annotation. Their relative numbers were compared across imputation tools.

## Abbreviations

log2-FC: log-2 Fold Change
MCC: Matthew’s Correlation Coefficient
MG: Müller Glia
MSC: Mean Silhouette Coefficient
scRNA-seq: single-cell ribonucleic acid sequencing
SD: Standard Deviation
TOM: Topological Overlap Matrix
t-SNE: t-distributed Stochastic Neighbor Embedding
WGCNA: Weighted Gene Correlation Network Analysis

## -Declarations-

### Ethics approval and consent to participate

No ethics approval was required for the study.

### Author’s contribution

LS, SC and JH designed the study. LS carried out the implementation, analysed the data and performed the calculations. LS, SC and JH wrote the manuscript. All authors have read and approved the manuscript.

### Availability of data and materials

The R-markdown files including reference data generation, data imputation, module preservation and edge recovery as well as the jupyter notbook including the marker gene analysis are provided on github (https://github.com/yigbt/scImpNetworks). Data used for downsampling can be found on https://github.com/software/github/SCRABBLEPAPER and the human retina organoid dataset at GEO:GSE119343. The gold data as well as corresponding dropout datasets are included in the R-package scorrgoldnet on github (https://github.com/lisbeth-dot-95/scorrgoldnet).

### Consent for publication

Not Applicable.

## Acknowledgements

Not Applicable.

## Footnotes

1 https://www.scrna-tools.org/analysis

2 https://github.com/yigbt/scImpNetworks

3 https://github.com/software-github/scrabble_paper

4 https://github.com/yanailab/enhance-R

5 http://loompy.org/

6 https://nbviewer.jupyter.org/github/iyhaoo/DISC/blob/master/reproducibility/Data\%20Preparation\%2C\%20Imputation\%20and\%20Computational\%20Resource\%20Evaluation/Data\%20Pre-processing/MELANOMA.ipynb

7 https://horvath.genetics.ucla.edu/html/CoexpressionNetwork/Rpackages/WGCNA/Tutorials/

