## Supplemental Material for "Benchmarking scRNA-seq imputation tools with respect to network inference highlights deficits in performance at high levels of sparsity"

### S1 Supplement

Table S1: Results of retina annotation pipeline before and after imputation. The table shows the percentages of retina specific cell types as well as not assignable and mixed clusters.

| | MSC | Clusters | Cones | Rods | MG cells | Not assignable | Mixed annotation |
|---|---|---|---|---|---|---|---|
| sparse | -0,01 | 8 | 26,03 | 21,60 | 24,34 | 28,04 | 0,00 |
| DrImpute | -0,18 | 11 | 24,29 | 21,10 | 18,05 | 6,55 | 0,00 |
| SAVER | -0,03 | 8 | 26,08 | 18,05 | 19,76 | 36,11 | 0,00 |
| DCA | 0,14 | 8 | 28,75 | 20,28 | 29,12 | 21,84 | 0,00 |
| scNPF | -0,03 | 9 | 20,43 | 21,77 | 23,25 | 34,55 | 0,00 |
| scNPF String | -0,02 | 8 | 20,65 | 21,03 | 18,50 | 39,82 | 0,00 |
| ENHANCE | 0,02 | 21 | 44,28 | 17,09 | 15,82 | 19,32 | 3,49 |

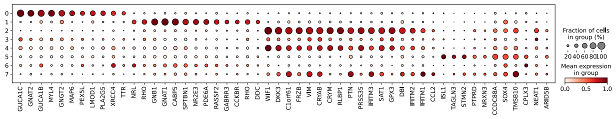

(a) Sparse human retina organoids

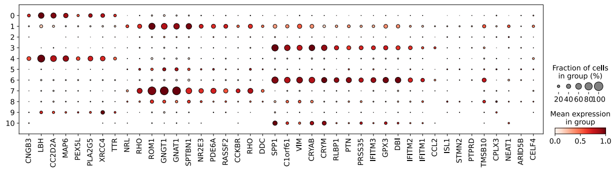

(b) `DrImputed` imputed human retina organoids

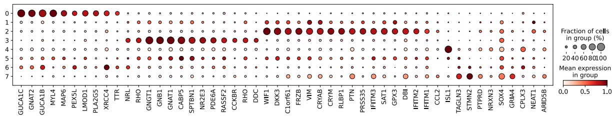

(c) `SAVER` imputed human retina organoids

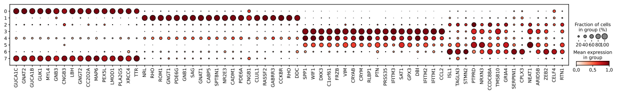

(d) `DCA` imputed human retina organoids

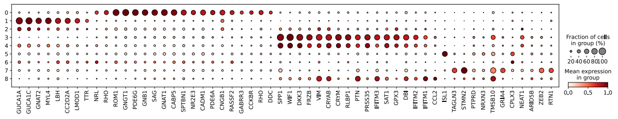

(e) `scNPF` imputed human retina organoids

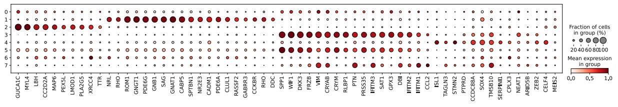

(f) `scNPF String` imputed human retina organoids

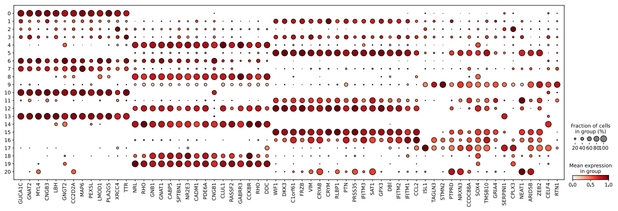

(g) `ENHANCE` imputed human retina organoids

Figure S1: Dotplot of marker gene expression in human retina organoids.
All dotplots show the expression values of marker genes across Louvain clusters using the `scanpy` workflow before and after imputation. Coloring corresponds to the mean gene expression and the dot size to the percentage of cells per cluster expressing the respective gene. Expression values were scaled per Louvain cluster.

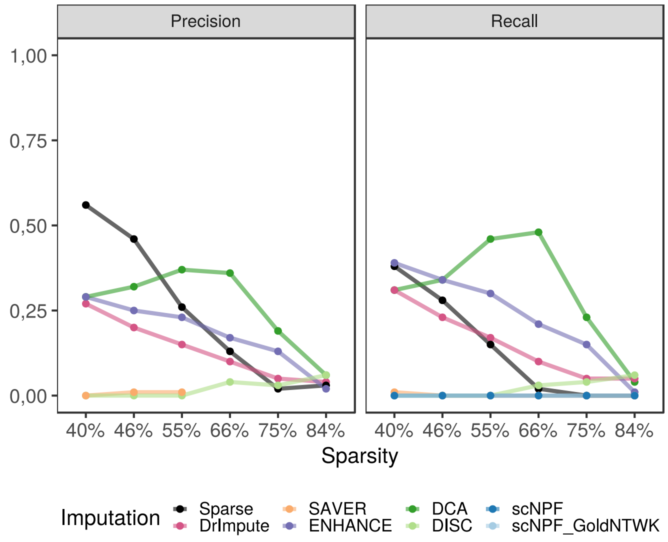


(a) Precision of edge recovery of second threshold

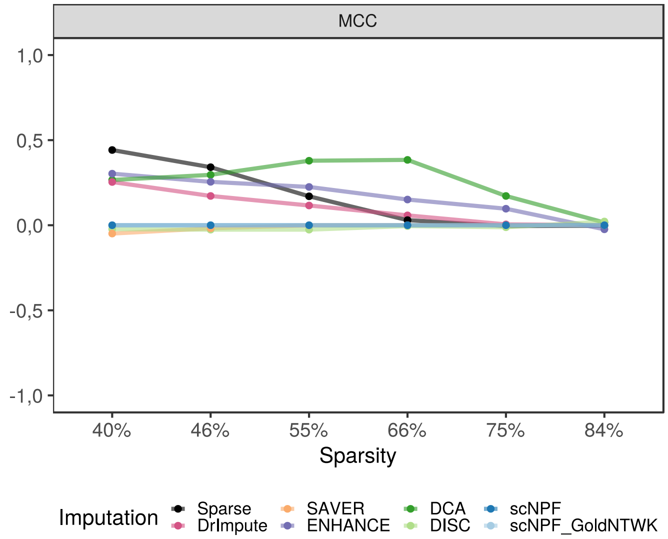


(b) Recall of edge recovery of second threshold

Figure S2: Additional threshold edge recovery analysis.
Trends of Precision, Recall (a), and Matthews correlation coefficient (MCC) (b) over all dropout levels indicating the ability to recover true edges after binarizing the gold data network. Mean and standard deviation ($SD$) of the topological overlap matrix (TOM) per dataset were computed. Edges with a TOM greater $2 * SD + mean$ of the gold and all dropped out and imputed datasets were retained. Retained edges of gold data were considered as true edges for computing Precision, Recall, and MCC, respectively.

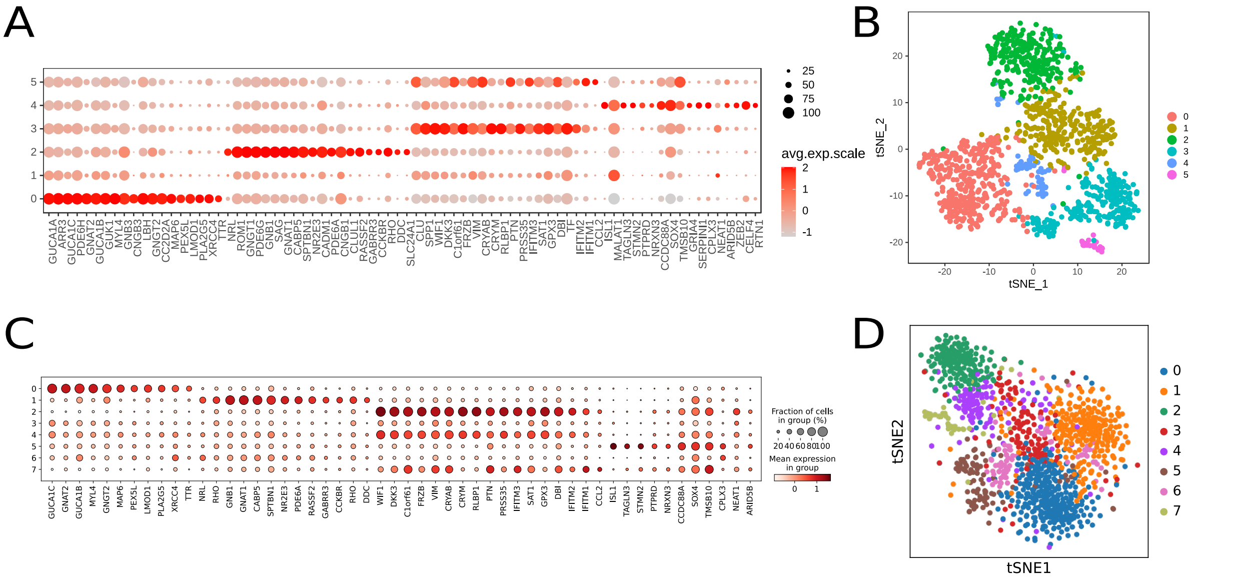


Figure S3: Comparison of `Seurat` and `scanpy` preprocessing results.
(A) Dotplot showing the expression of marker genes across Louvain clusters using the `Seurat` pipeline. Coloring corresponds to the mean gene expression and the dot size to the percentage of cells per cluster expressing the respective gene. (B) t-SNE of Louvain clusters using `Seurat`. (C) Dotplot showing the expression of marker genes across Louvain clusters using the `scanpy` pipeline. The expression was scaled per Louvain cluster. (D) t-SNE of Louvain clusters using `scanpy`. Colors between t-SNE plots do not correspond.

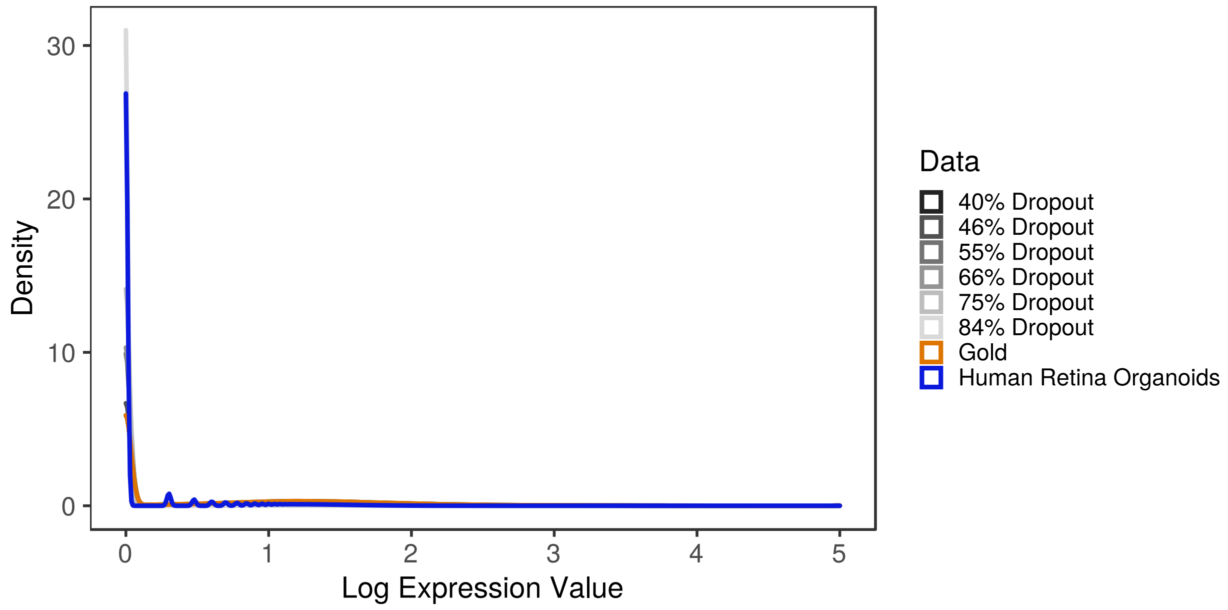


Figure S4: Distribution of logged expression values of all reference datasets and the human retina dataset.
Density of expression values across eight datasets is shown to contrast impact of different dropout rates. The gold data is plotted in orange, all six dropout reference datasets are shown with a grey gradient and a biological dataset is plotted in blue.

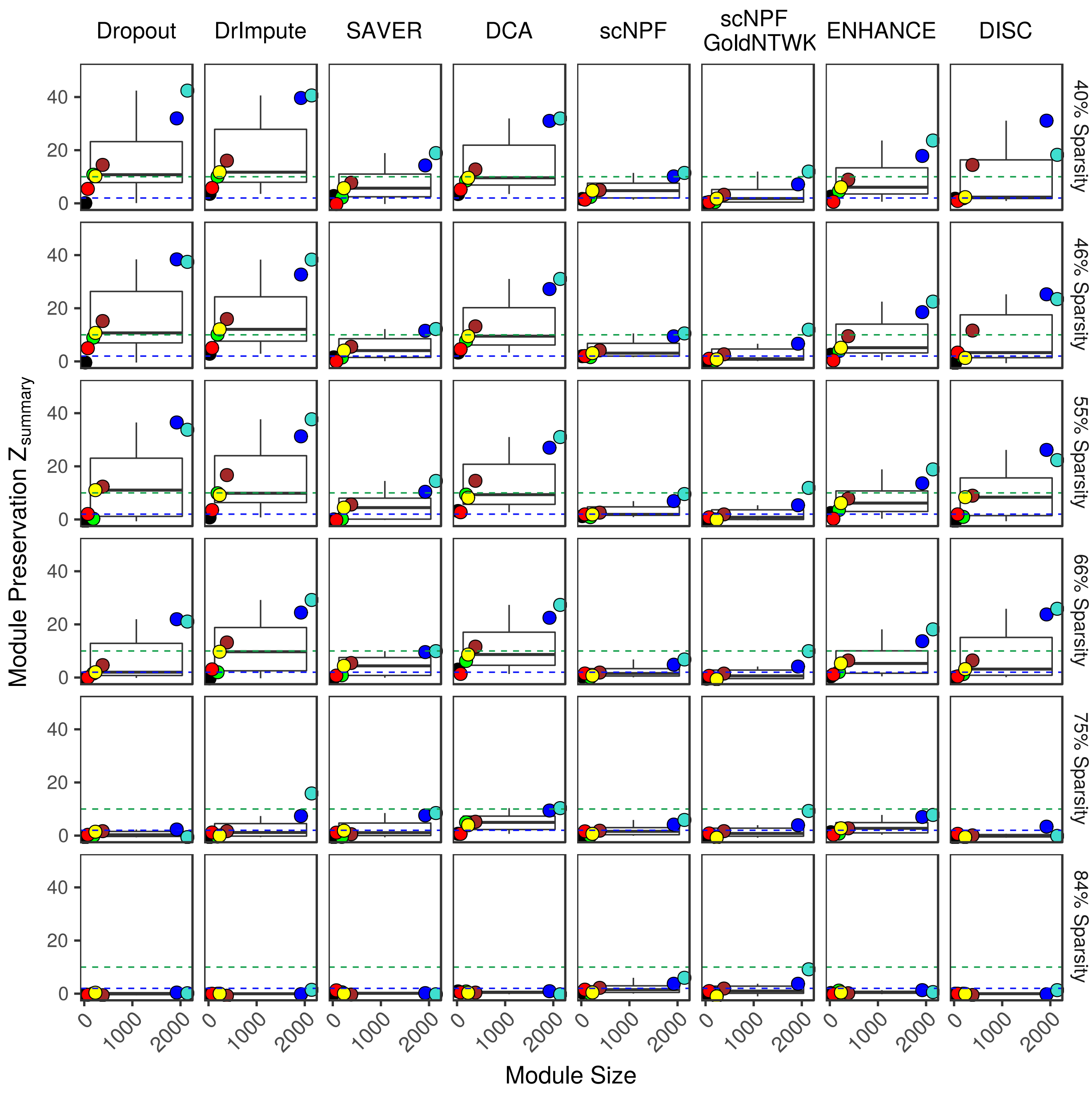


Figure S5: Boxplots showing the behavior of module preservation across different dropout levels. The $Z_{summary}$ measure implemented in `WGCNA` is a composite, permutation- based metric of various network density and connectivity measures. The blue and the green line indicate the threshold towards moderate and strong module preservation, respectively. Coloration of the dots correspond to the individual modules. Dropout refers to the the amount of artificially introduced non-true zeros in each of the reference datasets.

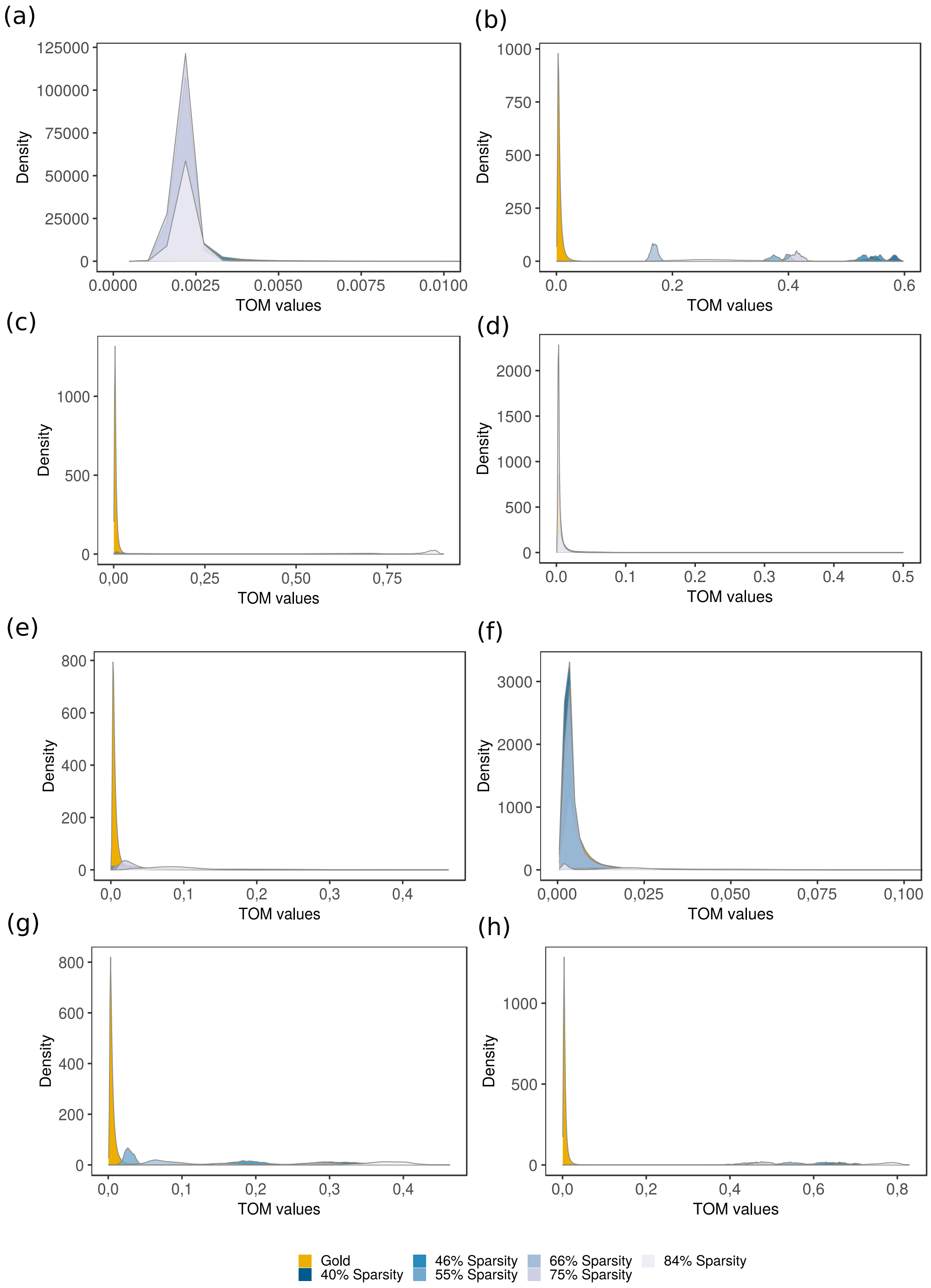


Figure S6: Density of TOM values before and after imputation of the synthetic data set. After transforming expression data before and after imputation to TOM values via the gold data $\beta$ value, positive edges were detected and compared during the edge recovery analysis. Colours correspond to original information content of data (lightest blue - less information, highest sparsity). (a) Dropout data, (b) `DrImpute`, (c) `SAVER`, (d) `ENHANCE`, (e) `DCA`, (f) `DISC`, (g) `scNPF`, (h) `scNPF Gold`. Whereby some tool, such as `DrImpute` (b) and both `scNPF` approaches (g+h) produce high TOM values compared to the gold data, `DCA` (e) and `ENHANCE` (d) TOM densities were close to gold.
